# Fertilization impacts microbiomes along the grassland trophic chain

**DOI:** 10.1101/2024.12.06.627205

**Authors:** Karoline Jetter, Kunal Jani, Kerstin Wilhelm, Ulrike Stehle, Rostand Chamedjeu, Christian Riedel, Lena Wilfert, Patrick Schäfer, Simone Sommer

**Author notes:** These authors contributed equally (Karoline Jetter, Kunal Jani). Senior author (Christian Riedel, Lena Wilfert, Patrick Schäfer, Simone Sommer).

## Abstract

Agricultural grasslands are often managed intensively, influencing soil properties and microbial communities. This can result in significant community changes and challenges to health and function at different levels along the trophic chain. This study investigates how fertilization affects microbial communities in grassland ecosystems in multiple connected trophic compartments. Shifts in microbial composition occurred in response to fertilization with effects being host-dependent and changes being more pronounced in belowground compartments. Strong interactions between trophic levels facilitated transmission of bacterial genera from soil and roots to higher trophic levels. Pig slurry- derived microbes were found in all compartments, but their low prevalence suggests an indirect effect of fertilisation, primarily due to changes in nutrient availability. Our findings emphasize the importance of considering both individual compartments and trophic interactions in ecosystems to fully understand the effects of anthropogenic disturbances on environmental and human health.

## Introduction

On a global scale, intensive land use is a major driver of ecosystem function and biodiversity loss ^1,2^. Apart from croplands, grasslands are the main component of agricultural landscapes in temperate regions, providing forage for grazers such as cattle and sheep. To increase the yield of hay and silage for livestock feeding, livestock slurry or manure are widely used fertilizers in grassland ecosystems. These organic fertilizers improve soil nutrient content and resource availability, thereby increasing plant productivity ^3^. However, organic fertilizers do not only directly affect soil properties such as composition of nutrients or pH, but also add livestock-associated microbial communities ^4,5^. The microbial influx may displace natural members of the microbiota and introduce additional microbes with unwanted traits e.g. antimicrobial-resistant genes (ARGs) ^6-8^. Such effects may not be limited to soil-associated communities, but may also potentially be transferred along the trophic chain, from microbiomes to macrofauna, and their associated ecosystem functions ^9,10^.

There is growing evidence that intensification of land use and fertilization affects bacterial and fungal communities in soil, roots, as well as flowers and leaves, reducing the complexity of microbiome networks and causing microbial communities to become more homogeneous ^11-14^. The diversity and stability of host-associated microbial communities, such as root-associated microbiomes in plants and gut microbiomes in animals, are crucial for host health ^15-17^. These diverse microbial communities serve important functions in e.g. nutrient allocation and cycling, and pathogen defense ^7,15,18,19^ or act as a barrier to the accumulation of antimicrobial resistances in the environment ^20^. An exogenous disturbance of the microbiome might result in shifts in microbial composition and diversity leading to dysbiosis, with a loss of commensal bacteria and increase in pathogens within the microbiome, with potentially serious consequences for ecosystem function and host health. Therefore, based on the eco- holobiont framework and the One Health concept, it is important not to consider specific compartments of an ecosystem in isolation, but to assess the impact of environmental perturbations, such as fertilization, across abiotic and biotic levels and trophic compartments ^7,10,21,22^.

Although aboveground and belowground compartments may be differentially affected by land use ^1,23^, they are strongly interlinked at the community level and directly connected at the local scale through plants ^24^. Soil microbial communities redistribute and provide nutrients by decomposing plant litter and other soil organic matter, promote plant growth, and support plant stress resilience, making this soil-plant feedback essential for primary production and ecosystem functioning in grassland ecosystems ^25,26^. On the other hand, plants influence soil microbial communities by providing resources and selectively recruiting beneficial soil microorganisms via rhizodeposition into the rhizosphere (i.e. the soil immediately adjacent to plant roots) ^17,27,28^. Various detritivorous and herbivorous animal species feed on belowground soil organic matter, roots and plant litter, as well as aboveground plant biomass, thus ingesting soil- and plant-associated microbes into their gut ^29,30^. Similar to soil and the rhizosphere, these (macro)organisms are also affected by fertilization-driven disturbances and may serve as reservoirs and vectors for zoonotic pathogens and antibiotic resistant bacteria ^8,19,31^. Aboveground, flowers can act as reservoirs for microbes ^32^. As different pollinating insects share floral resources, flowers function as transmission sites for microbes in pollinators, making microbe transfer between flowers and pollinators bidirectional ^33,34^. These trophic interactions, and the importance of stable and balanced microbial communities for host and environmental health, reinforce the need to collectively study microbial communities associated with multiple links in the trophic chain to understand the impact of fertilizers on environmental health and ecological function. However, so far, studies examining the effects of land use and fertilizers on microbial species richness and diversity have largely focused on isolated compartments in complex ecosystems ^1,23,35^.

In this study, we investigated the effects of four fertilization regimes on microbial diversity, composition, and interconnectedness across seven compartments along the trophic chain in grassland ecosystems. Specifically, we aimed at addressing the following questions: 1) Does fertilization affect microbial diversity similarly in above- and belowground, as well as in animal- and plant-related compartments of the trophic chain? 2) Are changes in microbial communities driven by a population of commensal allochthonous bacteria, or are they due to indirect effects affecting the indigenous microbial community? And 3) to identify microbial taxa that are shared and potentially exchanged between different trophic compartments? We postulated that belowground microbial communities would be strongly affected by the application of fertilizer. We hypothesized that pig slurry fertilization is the most intensive fertilization regime in our set up, and would therefore have the strongest effects through the introduction of allochthonous bacteria. Our findings confirmed that fertilization significantly affected the diversity of belowground compartments and the microbial composition of all trophic compartments, with pig slurry having the strongest effects. However, rather than direct effects through the establishment of allochthonous taxa, we observed indirect effects driven by changes in soil nutrient content. The effects of fertilization were strongly host-dependent, highlighting the need to assess the impact of fertilization across trophic compartments within a One Health perspective.

## Materials and methods

### Study area and sample collection

Sample collection was carried out between May and July 2022 on agricultural grassland sites located on the Swabian Alb in Baden-Württemberg, Germany (see Figure S1 (A)) that were subjected to four different fertilization regimes typical for this region i.e. fertilization with (i) biogas digestate, (ii) cow/horse manure, (iii) pig slurry and (iv) control sites, i.e. former military sites that are minimally managed by grazing flocks of sheep (maximum twice per year). In total, 28 grassland sites were investigated, including seven control, eight biogas digestate, seven cow/horse manure and six pig slurry fertilized sites. Sampling specimens were selected by targeting organisms that are representative of different trophic compartments above and belowground (Figure S1(B)) containing animals (zoosphere; i.e. feces of earthworms and voles (Microtus arvalis, Arvicola amphibious), bumblebee gut contents (Bombus lapidarius)) and plant-related samples (phytosphere; i.e. endosphere, rhizosphere and flowers of Trifolium pratense), as well as soil and samples from pig slurry. T. pratense L. is an important legume forage crop in temperate regions due to its high protein content^36^ and the symbiosis with nitrogen fixing bacteria ^37^. As T. pratense flowers are frequently visited by bumblebees and honeybees 38 and as several vole species prefer T. pratense roots as a food source ^39–41^, this system links above- and below-ground compartments in a widely distributed trophic chain in grassland ecosystems. In addition to the analysis of the microbial communities, the nutrient content of the soil was also analyzed. Details of sampling and initial sample processing are provided in the supplement.

### DNA extraction, library preparation and 16S rRNA gene amplicon sequencing

A total of 494 samples were collected. Details on sample sizes per compartment are given in Table S1. To assess the bacterial composition, the V4 hypervariable region of the 16S rRNA gene was amplified in a two-step polymerase-chain reaction (PCR) using the primers 515 F (5′-GTGCCAGCMGCCGCGGTAA- 3′) and 806 R (5′-GGACTACHVGGGTWTCTAAT-3′) ^42^. To avoid amplifying high amounts of chloroplast and mitochondrial DNA, peptide nucleic acid (PNA)-DNA clamps (mPNA and pPNA for plant (endosphere, flower) and mPNAs (bumblebee) samples) were added to the first PCR reaction ^43^. Paired- end sequencing of the constructed libraries was conducted in two separate runs on an in-house Illumina MiSeq sequencing platform according to Fackelmann et al. ^44^ and Heni et al. ^45^ (see supplement for details). DNA extraction and sequencing of a total of 494 samples (excluding negative and positive control) resulted in 10,044,014 reads and 27,753 ASVs after filtering.

### Soil nutrient assessment by electro-ultrafiltration

Soil samples were pooled from each sampling site, resulting in 23 pooled samples, and sent to the Bodengesundheitsdienst GmbH (https://www.bodengesundheitsdienst.de) for nutrient content analysis. Using electro-ultrafiltration (EUF), the concentrations of macronutrients (nitrate, organic nitrogen, phosphorus, potassium, sulfur, boron, magnesium, and calcium), micronutrients (iron, manganese, zinc, and copper), and sodium were determined for each site. EUF separates nutrients into two fractions based on different water pressures and membrane types: the first fraction represents nutrients immediately available to plants, and the second contains nutrients that are deliverable over time. For our analysis, we combined both the immediately available and deliverable fractions to calculate the total amount of nutrients for each site. We used the information of the content of eight macronutrients (P, K, S, N, nitrate, Ca, Mg, B) and four micronutrients (Mn, Zn, Cu, Fe) as well as the sodium content in soil samples for subsequent analyses.

### Bioinformatic sequence processing and statistical analyses

Sequence processing was conducted using the DADA2 pipeline in QIIME2 (version 2022.8) ^46^. This included the removal of primers, denoising, removal of chimeras and merging of paired-end reads ^47^. Further details can be found in the supplementary material. All statistical analyses were performed using the R programming language and the R Studio integrated development environment ^48^. In order to evaluate the impact of fertilization on the microbiomes of all seven different trophic compartments, measures for microbial alpha diversity (Shannon index) and beta diversity (Bray-Curtis distances) were calculated for each compartment separately. The differences between the fertilization regimes within each compartment were evaluated using analysis of variance (ANOVA) with Tukey’s HSD post-hoc test for alpha diversity and pairwise PERMANOVA (999 permutations) for beta diversity. The latter was conducted using the vegan package ^49^ and the pairwise Adonis package ^50^.

To determine the extent to which different soil nutrients drive differences in soil microbial composition, a redundancy analysis (RDA) was performed using Bray-Curtis distances of the soil microbial community and the available nutrients (P, K, S, N, NO3, Na, Ca, Mg, B, Mn, Zn, Cu, Fe) as constraints. The ordiR2step function implemented in the vegan package was employed to initially construct a forward selection model with the objective of identifying the nutrients that significantly explained the observed variation in microbial composition between the different soil samples.

Subsequently, the ordinate function, implemented in the microviz package ^51^, was utilized to run the RDA models separately for macronutrients, micronutrients, and all nutrients individually. This was done to determine the extent to which the nutrients in question could explain the variation between microbial communities.

Differentially abundant taxa, i.e., taxa that respond to fertilization within the different trophic compartments, were identified using Analysis of Composition of Microbiomes (ANCOMBC2) ^52,53^. In the first step, samples from all three fertilization regimes were compared individually against the samples from the control sites. In a second step, samples from all fertilization regimes were compared against each other (all pairwise comparisons). Core microbiome analyses were conducted for each trophic compartment individually, encompassing all pairwise comparisons between fertilization regimes to determine resilient taxa, i.e. taxa present in all samples within a compartment regardless of the fertilization regime, utilizing the core function in the microbiome package ^54^.

Subsequent analyses were conducted on a refined dataset, comprising only those ASVs identified as responding taxa in the differential abundance analysis and as resilient taxa in the core microbiome analysis. To verify the impacts of fertilizer application on microbial composition within the various trophic compartments, the refined dataset was employed to conduct a canonical analysis of principal coordinates (CAP) ^55^, followed by PERMANOVA and pairwise PERMANOVA. Based on interactions among trophic compartments within the trophic chain, a Source Tracker2 ^56^ analysis was conducted to assess microbial taxa that overlap between compartments and to elucidate potential exchanges of taxa between compartments. All graphs were created in R using the ggplot2 package ^57^.

## Results

### Impact of fertilization on microbial alpha and beta diversity in different compartments along the trophic chain

In summer 2022, a total of 494 samples from seven different trophic compartments were collected from agricultural grassland sites on the Swabian Alb (Germany) under four different fertilization regimes: control sites, sites fertilized with cow and horse manure, sites fertilized with biogas digestate and sites fertilized with pig slurry. Samples were divided into the plant-related “phytosphere” and the animal-related “zoosphere”, and we further distinguished between “aboveground” and “belowground” compartments of the trophic chain (Figure 1B).

**Figure 1:**
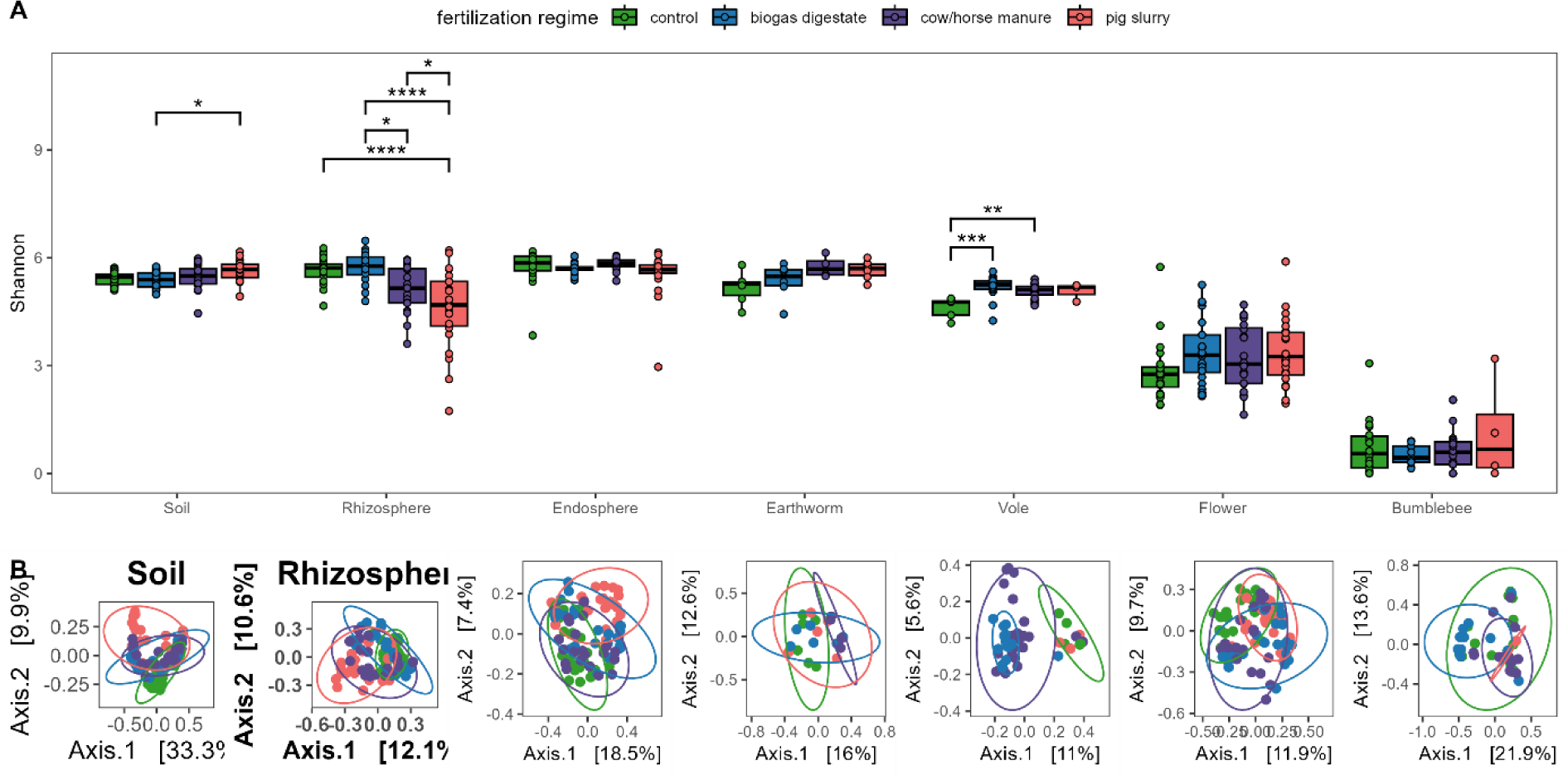
Effects of fertilization on microbial diversity. (A) Alpha diversity (Shannon index) and (B) Beta diversity (Bray Curtis) measures for individual compartments along the trophic chain. Green: control, blue: biogas digestate, purple: cow/horse manure, pink: pig slurry. ANOVA followed by post hoc Tukey’s HSD test was used to analyze effects of fertilization regimes on alpha diversity (*=p=0.05, ** p<0.01, *** p<0.001 **** p<0.0001). Differences in beta diversity were assessed using PERMANOVA followed by pairwise PERMANOVA.

Both microbial alpha (ANOVA: Shannon*, p*<0.001) and beta diversity (PERMANOVA: Bray-Curtis, df=3; R^2^=0.027, *p*=0.001) were highly variable between trophic compartments and fertilization regimes (Figure 1). Disentangling the overall significant effect of fertilizer application on alpha diversity revealed that Shannon diversity was significantly impacted by fertilization in the belowground compartments soil (ANOVA: *p*=0.029) and rhizosphere (ANOVA: *p*<0.001) as well as in the belowground zoosphere compartment voles (ANOVA: *p*=0.001) (Figure 1A, Table S2). In soil samples, alpha diversity was increased in pig slurry fertilized sites when compared to biogas but not to control or cow/horse manure fertilized sites (TukeyHSD: biogas vs. pig: *p*=0.029). For voles, alpha diversity was increased in biogas- and cow/horse-manure-fertilized sites (TukeyHSD: control vs. biogas: *p*<0.001; control vs. cow/horse manure: *p*=0.003). Moreover, alpha diversity of the rhizosphere was significantly lower in pig slurry-fertilized sites as compared to all other treatment groups (TukeyHSD: control vs. pig: *p*<0.001; biogas vs. pig: *p*<0.001, cow/horse manure vs. pig*: p*=0.049; biogas vs. cow/horse manure: *p=*0.026) (Table S2). Flowers and bumblebees, representing the aboveground trophic compartments, as well as earthworms exhibited no differences in alpha diversity (Figure 1A). Similarly, the root endosphere showed no significant variation under the different fertilization regimes (Table S2). Hence, the impact of fertilizer application is more pronounced in belowground compartments.

Beta diversity was significantly affected by fertilization regimes in all trophic compartments (Figure 2B). Within the phytosphere and soil compartments, we noted significant differences in beta diversity revealing an impact of all four fertilization regimes on microbiome compositions (soil: *p*=0.001, rhizosphere: *p*=0.001, endosphere: *p*=0.001, flower: *p*=0.001; PERMANOVA: Bray-Curtis; Table S3). Here, microbial communities differed between all four fertilization regimes, except for soil, where biogas and cow/horse manure fertilized sites did not differ significantly from each other (pairwise PERMANOVA: *p*=0.153). Similarly, significant differences in beta diversity between fertilization regimes were observed in all zoosphere compartments (voles: *p*=0.001; earthworms: *p*=0.005; bumblebees: *p*=0.001 PERMANOVA: Bray-Curtis; Table S3). In voles, microbial composition differed between all fertilization regimes except for comparisons between control and pig slurry fertilized sites (pairwise PERMANOVA: *p*=0.656). Pig slurry and cow/horse manure fertilization altered earthworm microbiomes (pairwise PERMANOVA: control vs. pig: *p*=0.021; control vs. cow: *p*=0.002) and fertilization with biogas digestate impacted the microbial compositions in bumblebees (pairwise PERMANOVA: control vs. biogas: *p*=0.023; cow vs. biogas: *p*=0.001; pig vs. biogas: *p*=0.005) (Table S3). Overall, these analyses revealed an impact of fertilization on microbial communities of all trophic compartments and pig slurry appeared to have the most pronounced effects.

**Figure 2:**
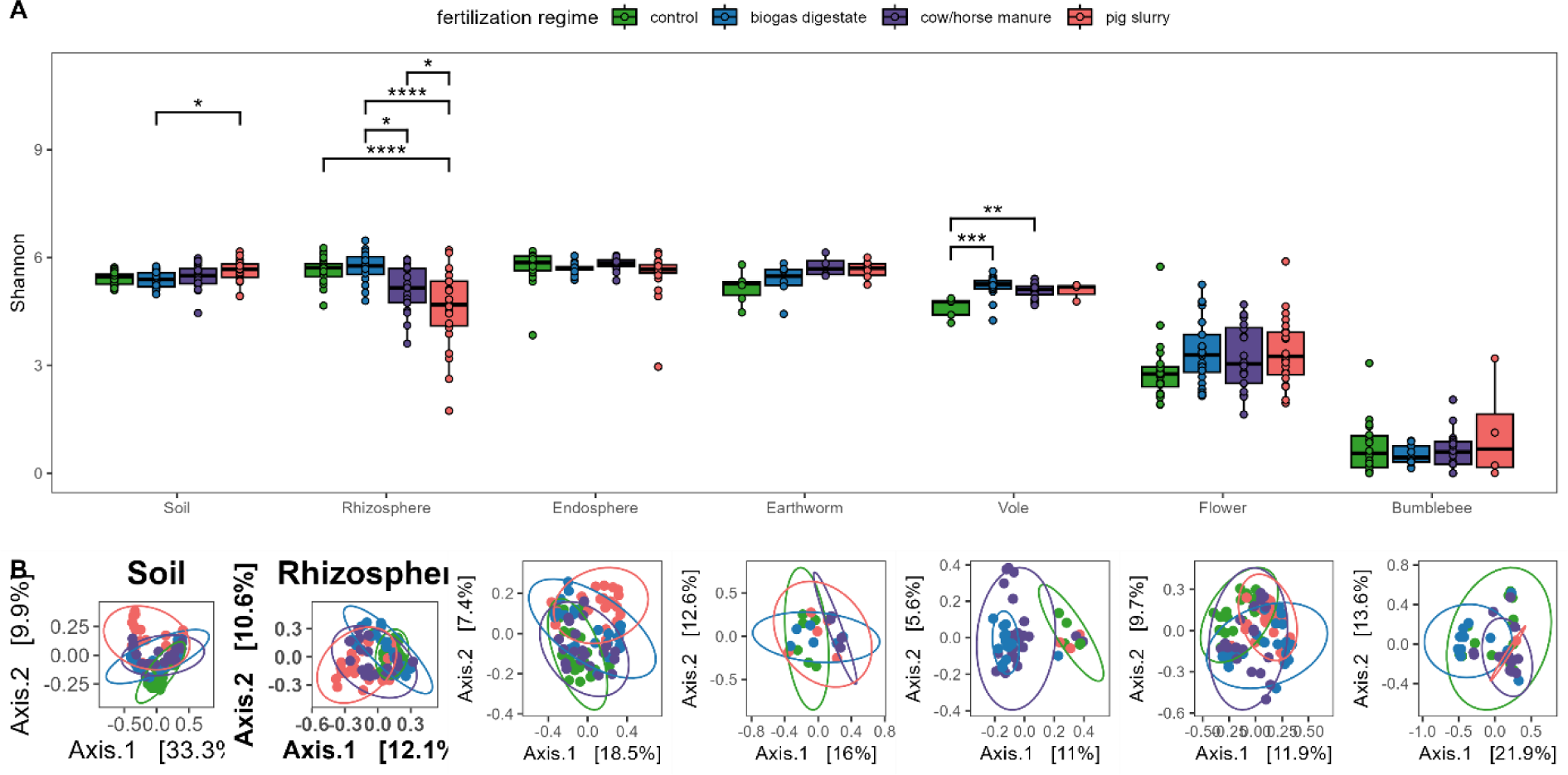
Bacterial composition of the different compartments along the trophic chain. Indicated are genera with a minimum abundance of 5%. Phytosphere compartments are marked in green, zoosphere compartments are marked in pink, soil is marked in brown. Data was pooled for plotting based on the fertilization regime within each compartment. Every fertilization regime contains up to 6 plots with 4 samples each, i.e. each bar represents a pool of up to 24 samples (see Table S1 for sample sizes).

### Effects of soil nutrients on soil microbial communities

Redundancy analysis (RDA) was used to model the amount of variation between soil microbial communities of different samples based on Bray-Curtis distances that can be explained by the different soil nutrients. Soil electro-ultrafiltration provided the content of eight macronutrients (P, K, S, N, nitrate, Ca, Mg, B) and four micronutrients (Mn, Zn, Cu, Fe) as well as the sodium content in soil samples. Forward selection of significant values revealed non-significant impacts of zinc within micronutrients and magnesium within macronutrients. After selecting significantly impacting nutrients, the model explained 51.9% of variation between the different soil microbial communities.

Selected macronutrients explained 38.9% of variation (adjusted R^2^=0.389; Figure S2A) while selected micronutrients including sodium explained 28.2% of variation (adjusted R^2^=0.282; Figure S2B) between microbial communities of different samples. Out of all tested nutrients, calcium explained the most variation (19.2%), followed by phosphorous (11.1%) and nitrate (9.1%) while only phosphorous contents were significantly different between soils collected on the four different fertilization regimes (ANOVA: *p=0.035*; Figure S2C).

### Fertilization affects microbial composition of different trophic compartments

There were substantial differences in how fertilization affected the microbial composition of the different compartments along the trophic chain at the phylum level. In phytosphere compartments, we detected lower abundances of *Firmicutes* in samples collected on fertilized sites as compared to control sites with strongest reduction caused by fertilization with pig slurry. Belowground compartments (soil, rhizosphere and endosphere) additionally differed in the abundance of *Verrucomicrobiota*. For the zoosphere compartments, *Firmicutes* were less abundant in fertilized plots for both earthworms and bumblebees, while microbial abundances in vole guts were stable at the phylum level across treatments (Figure S3A, Supplements).

The fertilizer effects on microbial composition were more prominent in all trophic compartments at the genus level. In samples of collected pig slurry, *Clostridium_sensu_stricto_1* and *Terrisporobacter* stood out, followed by *Pseudomonas*, *Turicibacter*, *Lactobacillus and Romboutsia* (Figure S3B). Across belowground compartments, the most prominent differences in fertilized sites compared to control sites were lower abundances in *Bacillus* and higher abundances in *Pseudomonas* (Figure 2). In soil, the predominant genera at control sites were *Bacillus* and *Rokubacteriales* whereas *Gaiella* was the dominant genus in fertilized sites. Although *Rokubacteriales* were one of the dominating genera in biogas fertilized and cow/horse manure fertilized as well as control sites, they were present only in lower abundances in pig slurry fertilized sites. Despite differences in abundance*, Pseudomonas* was the predominant genus in the rhizosphere in all fertilization regimes and in the endosphere in pig slurry fertilized sites. *Kineosporia,* which dominated the composition of the endosphere in control, biogas digestate and cow/horse manure fertilized sites, was far less abundant under pig slurry fertilization. *Flavobacterium* and *Bacillus* dominated the earthworm composition and *Solirubrobacter* were more abundant in fertilized sites as compared to control sites. Microbial composition in voles was dominated by *Ruminococcus* and two *Clostridia* groups in all treatments. *Gastranaerophilales* were present only in fertilized sites while *Roseburia* and *Acetatifactor* were less abundant in fertilized sites. In flower compartments, *Massillia* and *Ralstonia* were dominant in samples collected on biogas and pig slurry fertilized sites, while *Duganella,* which dominated samples from control sites, were less abundant in fertilized sites. In bumblebees, where *Gilliamella* were most abundant in all fertilization regimes, *Pseudomonas* were more abundant and *Lactobacillus* were less abundant on fertilized sites compared to control sites. Genera dominating pig slurry composition were detected in all the compartments across the trophic chain with relative abundances between 1% and 5%, so these analyses suggest an indirect effect of fertilization on the microbiome compositions of the different compartments.

### Fertilization impacts on resilient and responding bacterial taxa along trophic compartments

To obtain insights into which genera are significantly driving the changes in microbial communities between fertilization regimes, we analyzed the quantitative impact of fertilization on taxa of all trophic compartments collected at the different grassland sites using ANCOMBC2. This analysis detected 600 ASVs belonging to 82 genera (Figure 3A, Figure S4). As the abundance of these genera varied within trophic compartments in response to different fertilization regimes, we refer to these as responder taxa. In total, 41 taxa were increased whereas 25 taxa were decreased in abundance upon fertilizer application.

**Figure 3:**
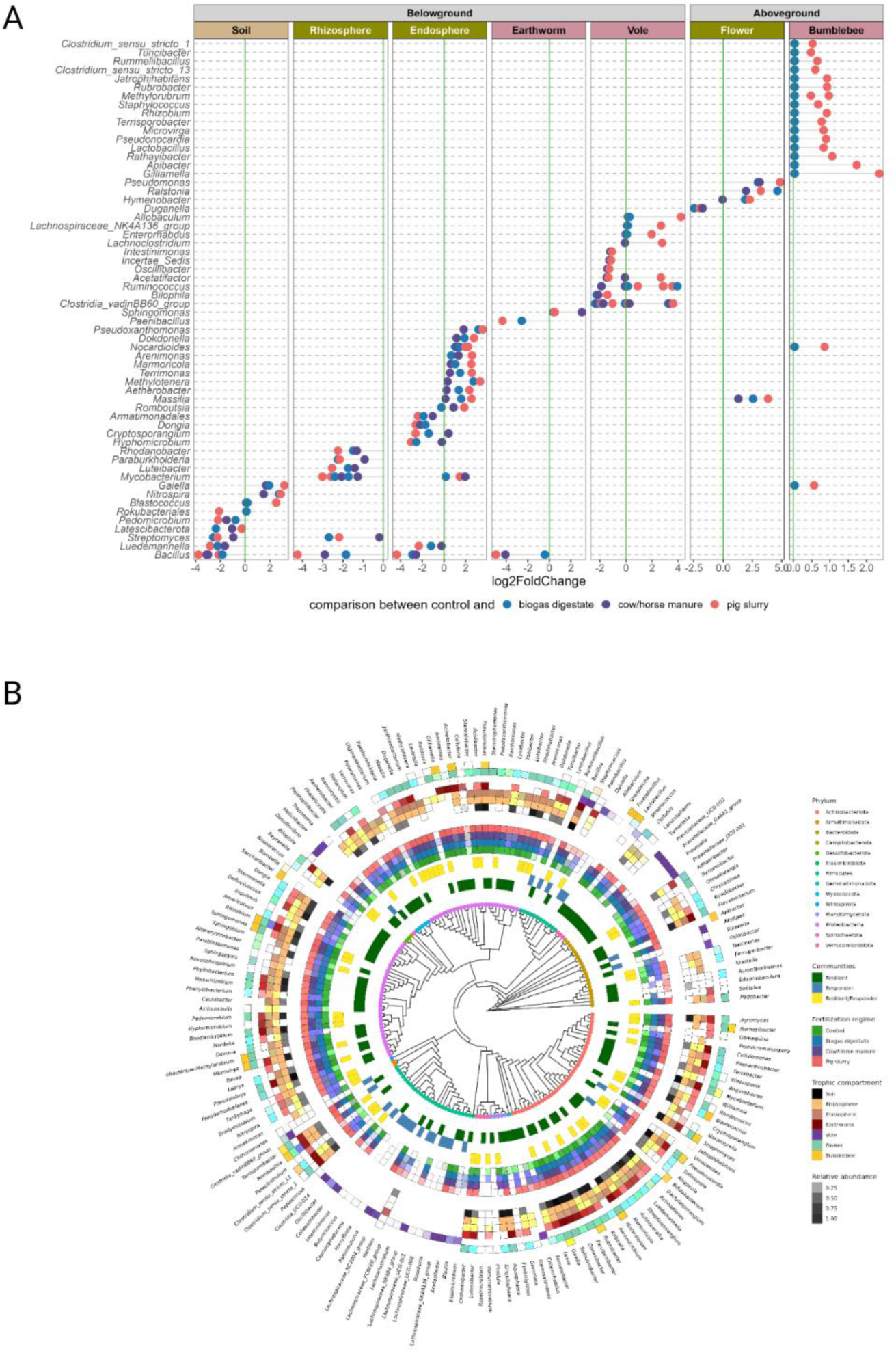
Bacterial genera and taxa responding to fertilization within the different trophic compartments. (A) Differentially abundant genera between unfertilized control sites (green line) and sites fertilized with biogas digestate (blue), cow/horse manure (purple) and pig slurry (pink), respectively, were identified using ANCOMBC2. Phytosphere compartments are marked in green, zoosphere compartments are marked in pink, soil is marked in brown. Positive values of log2fold change represent genera that are more abundant on fertilized sites; negative values represent genera that are less abundant on fertilized sites compared to control sites. (B) Responding (differentially abundant) and resilient taxa (members of the core microbiome) according to fertilization regimes and trophic compartments they were found in.

By running a core microbiome analysis on every individual compartment covering all pairwise comparisons of fertilization regimes, we determined the core microbiome of an individual compartment irrespective of the fertilization. Taking the results of the individual analyses of all seven compartments together, core microbiomes contained 894 ASVs associated to 159 identifiable genera across the whole dataset. Given that these genera are present within a trophic compartment across all fertilization regimes, we refer to them as resilient taxa (Figure S5). Comparing the genera of resilient and responding taxa across all trophic compartments, we identified 57 overlapping genera. These genera can be considered as fertilization-responsive core genera, due to their presence in all fertilization regimes within one trophic compartment and their fertilizer-dependent, differential abundance (Figure 3B).

In all phytosphere compartments, as well as in the soil and in earthworms, the abundance of resilient genera was higher in comparison to responder genera and all responder genera overlapped with the resilient genera. Hence, changes in the composition caused by fertilizer application were driven by changes in abundance of core taxa. In bumblebees and voles, numbers of resilient genera were lower in comparison to responder genera. Here, only a few differentially abundant genera overlapped with the core microbiome and changes in the composition caused by fertilizer application were driven by opportunistic non-core genera.

Amongst trophic compartments, seven differentially abundant genera overlapped between at least two different compartments. In several belowground and plant-associated compartments (soil, roots, earthworms), *Bacillus, Luedemannella*, and *Streptomyces* were found to be differentially abundant and lower in abundance in samples collected on fertilized sites compared to those collected on control sites. Comparing fertilized sites to control sites, *Gaiella, Massilla* and *Nocardioides* were more abundant in samples collected on fertilized sites than on control sites in soil and endosphere as well as in flowers and bumblebees (Figure 3A).

In line with microbial compositions (Figure 2 and Figure S3) within the different resilient microbiota, five genera were found to be core to several compartments of the trophic chain. *Bacillus, Microvirga, Pseudonorcardia* and *Gaiella* were core genera in all belowground and plant-associated trophic compartments (soil, rhizosphere, endosphere, worm gut and flower). Additionally, *Sphingomonas* was a core genus in rhizosphere, endosphere, earthworm gut, flower and bumblebee microbiomes. Voles had no overlapping core genera, hence showing a very host-specific core microbiome (Figure S5). Based on these analyses, *Bacillus* and *Gaiella* stood out as two genera being core to several compartments and being differentially abundant in those genera, hence they are members of the core microbiome that are affected by fertilization. *Bacillus* was less abundant on fertilized sites, i.e. suffering from fertilization, while *Gaiella* was more abundant on fertilized sites and thus profiting from fertilizer application.

### Identification of microbial taxa that are shared and potentially exchanged between different trophic compartments

In order to elucidate the potential exchange of microbial communities across the different compartments, SourceTracker2 analysis was performed on the refined data set containing resilient and responder taxa. Potential sources of microbiota (soil, rhizosphere, endosphere and flowers) were identified according to their position in the trophic chain. Soil was assigned as a source for the rhizosphere, flower, earthworm and vole compartments; rhizosphere was assigned as a source for endosphere, earthworm and vole and endosphere was assigned as a source for earthworm and vole, whereas flowers was the source for bumblebees. All interactions considered in the SourceTracker2 analysis are presented in Table S4. The analysis of microbial exchange along the phytosphere compartments (using soil, rhizosphere and endosphere as source) suggested the exchange of 57 genera between soil and rhizosphere, followed by 47 genera between rhizosphere and endosphere and 53 genera between soil and flower. In earthworms, numerous signals of microbe exchange were found between soil (53 genera) and rhizosphere (42 genera) as well as the endosphere (32 genera). By contrast, we found minimal signals of exchange of microbial genera of voles with soil (2 genera), rhizosphere (1 genus) and endosphere (1 genus). Similarly, no evidence for microbiota exchange was observed between flowers and bumblebees (Figure 4).

**Figure 4:**
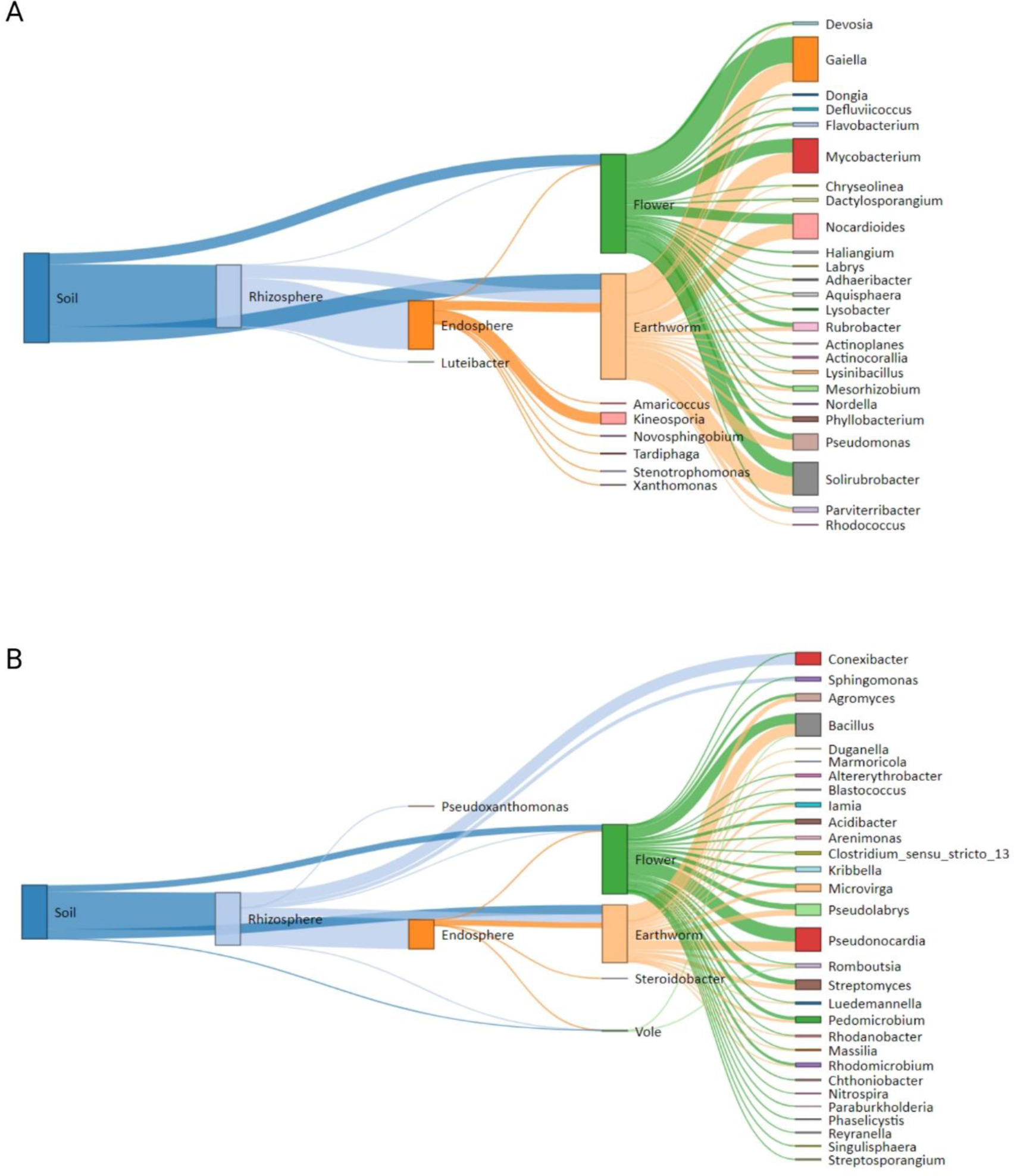
Proportions of bacterial genera shared between source and different target compartments along the trophic chain analyzed by SourceTracker2 analysis. A) Results based on genera identified as resilient taxa, B) Results based on genera identified as both, resilient and responding taxa. In both analyses, the source compartments were soil, rhizosphere and endosphere while key target compartments were endosphere, flowers, earthworms and voles.

Shared genera identified by the SourceTracker analysis were mostly taxa identified as resilient and responder, while taxa identified as only resilient were less frequently shared between compartments (Figure 4A-B). We found *Gaiella, Mycobacterium, Solirubrobacter, Nocardioides, Pseudomonas, Kineosporia, Rubrobacter, Mesorhizobium, Parviterribacter* and *Phyllobacterium*as to be the predominant resilient genera with evidence for exchange across different trophic compartments except for voles and bumblebees (Figure 4A). Similarly, *Pseudonocardia, Bacillus, Conexibacter, Pseudolabrys, Streptomyces, Agromyces, Microvirga, Pedomicrobium, Kribbella* and *Acidibacter* were found to be the most abundant genera identified as resilient and responder and were detected across different trophic compartments (Figure 4B). *Bacillus* and *Romboutsia* were present in all belowground compartments including voles. Concurrently, the selective exchange of *Rhizobium* and *Paenarthrobacter* was observed from the soil up to the rhizosphere and endosphere.

In line with the hypothesis that pig slurry fertilization is the most intensive fertilization, an examination of the exchange of microbes from pig slurry across trophic compartments was conducted. This revealed that four bacterial genera, namely *Clostridium sensu stricto 1, Terrisporobacter, Turicibacter* and *Romboutsia* were transferred from pig slurry to other compartments (Figure S6). The bacterial genus *Turicibacter* was detected in all trophic compartments up to bumblebee, whereas *Terrisporobacter* and *Romboutsia* were found in the flower compartment and all belowground compartments, except in voles.

Despite these shared genera across compartments and treatments, principle constrained analysis of principal coordinates (CAP) using the refined data set showed clear differentiation of the different trophic compartments (PERMANOVA: *p*=0.001, df=06, R^2^=0.389) and differences between the different fertilization regimes across all trophic compartments (PERMANOVA: *p*=0.001, df=3, R^2^=0.030) (Figure S7). Pairwise PERMANOVA revealed that all trophic compartments show a distinct microbial composition and differ significantly from each other in all pairwise compartment comparisons (Table S5). Further, pairwise PERMANOVA showed significant differences between all fertilization regimes across all trophic compartments (Table S6). Samples that have been collected from sites subjected to the same fertilization regime thus differ significantly from samples collected from all other fertilization regimes, irrespective of the trophic compartment.

## Discussion

Intensive land use, particularly activities like fertilization, can markedly affect grassland biodiversity at the microbial to fauna level. Fertilization can disturb the balance of microbial communities, which in turn can have far-reaching effects on the health of plants, animals, and entire ecosystems. Our study shows that fertilization alters the diversity and composition of microbes across different levels of the trophic chain in grasslands. We found that these changes in microbial communities can occur along trophic interactions suggesting that microbes are transferred between plants, soil, and animals. The strongest effects were seen in soil and plant-associated microbiomes. Our study emphasizes that microbiomes in ecosystems are interconnected, and understanding how external treatments like fertilization affect these microbiomes is crucial for protecting ecosystem and host health, as outlined in the One Health approach.

### Trophic compartment and type of organic fertilizer strongly influence the microbiome

Characterization of the microbiome across above- and belowground compartments along a grassland trophic chain revealed that fertilization increased microbial diversity in all compartments except the roots, specifically in the rhizosphere and endosphere. Here, the rhizosphere demonstrated a gradual yet significant decline in diversity, where alpha diversity decreased with fertilization intensity, from the highest diversity in the minimally fertilized control sites to biogas digestate, cow/horse manure and finally pig slurry fertilized sites. This reduced microbial diversity is linked to higher nutrient availability and associated reduction in plant-beneficial bacteria under high land use intensity ^3,58–60^. Indeed, fertilized sites in our study showed high levels of phosphorus and nitrate (NO3), with the highest levels found on pig slurry sites. Consistent with these results, reduced rhizospheric nitrification and amoA gene (ammonia monooxygenase) abundance were found in nutrient-enriched soils, particularly those treated with pig slurry and biochar ^61^. In contrast, voles showed a significant increase in gut microbial alpha diversity on biogas digestate and cow/horse manure sites. Additionally, contrasting effects of fertilization on aboveground and belowground trophic compartments were evident, with alpha diversity being strongly impacted in the belowground compartment, particularly by pig slurry. Le Provost et al.^1^ similarly demonstrated contrasting responses between aboveground and belowground trophic compartments by analyzing over 4,000 taxa, including soil microbes, from grasslands along a land-use gradient, including fertilization.

Fertilization also influenced microbial community composition, as demonstrated by clustering of microbial communities across all trophic compartments based on fertilization regimes. However, the effect of fertilization was highly dependent on the trophic compartment. As macro- and micronutrients account for 38.9% and 28.2% of the variation, respectively, these shifts highlight the impact of nutrient availability on microbial community structure emphasizing the indirect effects of fertilization. This supports the role of fertilization in shaping microbial communities in soil and roots, and in reducing complexity of the microbiome by altering soil properties along with the introduction of livestock-associated microbiota ^4,5,11,14^. These results are in line with those of Tyrrell et al.^62^, who investigated the impact of swine, bovine, and poultry manure on soil and grass phylosphere microbiomes, indicating deleterious effects of manure-based fertilizers on the microbiome.

It is therefore reasonable to assume that fertilization-induced alterations in soil and phytosphere microbiomes may also impact the microbiomes of dependent detritivorous and herbivorous animal species. This is in line with growing evidence of the longitudinal transfer of microbes from soil through roots to various plant shoot parts, including their transmission to pollinators via flower-pollinator interactions ^63–65^. Furthermore, trophic interactions and the recently proposed “eco-holobiont” framework ^21^ highlight the importance of considering both fertilization-mediated modulation and the potential exchange of microbes across trophic compartments ^24^. Under the eco-holobiont concept, the assembly of a holobiont within an organism is shaped by a circular microbial loop of interacting biotic and environmental microbiomes within an ecosystem. In other words, fertilization-induced perturbation of the microbiome, leading to dysbiosis, could trigger cascading effects throughout the trophic chain.

### Cascading effects of microbiota alterations across trophic chain

Using a Bayesian mixing model, we independently demonstrated the potential exchange of microbes across the trophic chain. We observed a unique assemblage of microbes categorized as resilient and resilient/responder across various trophic compartments, while the model suggested no exchange for responders. Resilient taxa, which represent core microbial communities consistently found across different fertilization regimes within each trophic compartment, were more prevalent than responder taxa, particularly in soil, the phytosphere, and belowground compartments, indicating their higher adaptability. Similarly, resilient/responder taxa—members of the core microbiome affected by fertilization—were widespread across compartments, however their characteristics remain largely unexplored. They may represent trophic compartment-associated commensals or pathobionts influenced by organic fertilizers. In contrast, responder taxa, which exhibited greater modulation in response to fertilization were significantly and consistently influenced by pig slurry. These responder taxa were particularly prevalent in zoosphere compartments, such as voles and bumblebees.

In light of the significant impact of pig slurry on microbiome composition, the assessment of potential exchange of prevalent pig slurry microbiota revealed presence of four genera—Clostridium sensu stricto 1, Terrisporobacter, Turicibacter, and Romboutsia—across multiple trophic compartments.

While it is often assumed that allochthonous communities displace indigenous communities ^4,5,61^, our findings reveal an indirect effect of these genera. Additionally, the abundance of pig slurry-associated taxa remained relatively low across all trophic compartments. However, their repeated detection across compartments indicates their potential to establish in these environments. Notably, Turicibacter includes three validly described species isolated from chicken eggshells, swine ileum, human blood, and mouse feces. The presence of these microbes across trophic compartments highlights the potential risk of livestock-associated microbes establishing themselves, thereby posing risks to both environmental and human health.

Fertilization-driven shifts in microbial communities, reduced complexity, and microbial influx could compromise host health. The affected host can serve as a reservoir for pathogens and microbes carrying antimicrobial-resistant genes (ARGs), thereby aiding their dissemination ^6–8^. For example, the established longitudinal transmission of microbes between soil and aboveground plant parts extends to flowers. Although microbial transfer between flowers and bumblebees was not observed, we did identify Turicibacter, a pig slurry-associated microbe, in bumblebees, as mentioned above. Since flowers are visited by multiple pollinators, they serve as both a reservoir of microbes and a transmission site, facilitating bidirectional dissemination while interacting with foraging pollinators, thereby affecting pollinator health ^32^. Such compromised hosts might amplify the risk of transmission of zoonotic pathogens ^63–65^. Likewise, below-ground animals that rely on organic matter (including roots and plant litter) also ingest soil and plant-associated microbes that affect their microbiome. These organisms are not exempt from microbial dysbiosis and can act as reservoirs or vectors for antibiotic-resistant bacteria and pathogens ^8,19,31^. The common vole is considered a pathogen reservoir and is known to play a significant role in the transmission of zoonotic diseases ^66^. Pathogen prevalence in Peromyscus mice has also been shown to increase in anthropogenically disturbed landscapes ^67^. Furthermore, rodents living close to human-built (disturbed) environments are likely to enhance the risk of zoonotic disease transmission, thereby posing a threat to human health ^66,68^. Moreover, the potential of fertilizer-driven spread of ARGs and mobile genetic elements (MGEs) in grasslands pose a grave concern. Breaking the cyclic transmission of ARGs and associated pathogens between animals and the environment has become increasingly difficult, which can have serious health implications in the context of One Health ^69^. However, these transmission dynamics are likely influenced by multiple factors and require interpretation in the context of specific hosts and the impact of introduced or displaced microbial communities, and thus merit further investigations.

In summary, land use change such as intensification of fertilization significantly modulates the microbiome across the trophic compartments of the grassland ecosystem. The differential responses of trophic compartments and the partial cascading effect of microbiome changes align with the eco- holobiont framework and may have serious implications for both animal and human health.

## Acknowledgement

We would like to thank our bachelor students Annika Eberl, Miriam Gillitzer, Juliana Jenke, Anna Jurtschenko, Sara Merico, Max Schorge and Tamara Sippert for their immense help with sampling and laboratory work, as well as Eva Köpff for her great help in contacting farmers. We thank the managers of the three Exploratories, Julia Bass and Max Müller, Anna K. Franke and Robert Künast as well as Franca Marian and Uta Schumacher and all former managers for their work in maintaining the plot and project infrastructure; Victoria Grießmeier for giving support through the central office, Andreas Ostrowski for managing the central data base. We thank the administration of the UNESCO Biosphere Reserve Swabian Alb as well as all land owners for the excellent collaboration. We further thank the four pig slurry farmers independent of the biodiversity Exploratories for their great collaboration and openness. This work has been funded by the Baden-Württemberg-Stiftung program “Mikrobiom” (ID 18 IMPALA, PI S. Sommer).

## Data availability

This work is based on data elaborated by the IMPALA project in cooperation with the Biodiversity Exploratories program (DFG Priority Program 1374). The datasets are publicly available in the Biodiversity Exploratories Information System (http://doi.org/10.17616/R32P9Q) and on NCBI (Bioproject number: PRJNA1188490). The datasets are listed in the reference section ^70,71^.

## Author Contributions

Conceptualization, S.S., L.W., C.R. and P.S.; Investigation, Ka.J., K.W., U.S. and R.C.; Formal Analysis, Ka.J. and Ku.J.; Data Curation Ka.J., Ku.J., S.S., L.W., C.R. and P.S.; Writing – Original Draft, Ka.J. and Ku.J.; Writing – Review & Editing, Ka.J., Ku.J., S.S., L.W., C.R. and P.S.; Funding Acquisition, S.S., L.W., C.R. and P.S.; Supervision, S.S., L.W., C.R. and P.S.

