## Supplementary material for "Fertilization impacts microbiomes along the grassland trophic chain": Jetter_Jani_supplementary.pdf

[5] Senior author

[6] Lead contact

[*] Corresponding author

### Supplementary methods

#### Sampling of focus specimens along the trophic chain

Soil, plant (red clover, *Trifolium pratense*), earthworm, bumblebee (*Bombus lapidarius*) and vole (*Microtus arvalis* and *Arvicola amphibius*) samples were collected in grassland ecosystems subjected to different fertilization regimes to investigate the effects of fertilization on microbiomes along the trophic chain (Figure S1B). All grassland sites except sites fertilized with pig slurry are part of the Biodiversity Exploratories framework (https://www.biodiversity-exploratories.de)[1]. The pig slurry fertilized sites were selected in cooperation with local farmers on the Swabian Alb (Figure S1B).

Soil, earthworms and plants were sampled together in an area of approximately 10cm x 10cm around a red clover stem. Soil was sampled by collecting the top layer (to a depth of 15 cm) from the 10x10x15 cm ditch and sifting through a 0.1 cm kitchen sieve to remove larger stones and roots. Samples of red clover *Trifolium pratense* were selected by visual inspection, choosing a healthy individual with at least one open flower at anthesis; large plants, usually characterized by large root systems, were excluded to facilitate root processing. Plant samples were collected by removing the plant with its roots from the ditch in order to retain the plant roots by carefully removing the bulk soil adhering to the roots. Plant height and number of flowers per plant were also recorded. Roots and flowers were separated from the aboveground biomass and all samples were kept on ice during transport to the laboratory for further processing. Due to the extremely dry climatic conditions during the summer of 2022, all earthworms were collected from the same 10x10x15cm ditch regardless of taxonomic affiliation. Earthworm feces were collected by placing the earthworms in a petri dish with a damp paper towel overnight and then releasing the earthworms. Bumblebees of the species *Bombus lapidarius* were captured at the field sites using a plastic container when visiting a flower inside or close to the site. Voles were captured using Sherman live traps with a bait containing a mixture of maltol solution, oat flakes and cereals, and an apple slice. The traps were modified according to Mori et al.[2] by attaching half a Tetra Pack carton with an entrance hole to the trap opening, which was then placed on a vole burrow. Traps were set in the late evening and checked early the next morning. If a vole was caught, its feces were collected in 400µl nucleic acid preservation (NAP) buffer [3] and the voles were released after recording their weight and sex. All samples were transported to the laboratory on ice for further processing and stored at -20°C until further analysis.

**Sample processing**

Plant samples were separated into root, flower and aboveground biomass samples. Root samples were separated into endosphere and rhizosphere samples (see Figure S1B). The rhizosphere was washed from the roots by shaking the whole root in 25 ml sterile water, removing the roots and centrifuging the solution at 4000 × g for 15 min to obtain microbial cells. Endosphere samples were obtained by shaking the root in 30 ml of 0.1% (v/v) Triton X-100 in $H_2O$ followed by washing off the Triton solution with 30 ml of sterile water. Roots were then air dried on sterile filter paper and cut into smaller pieces for extraction. Flowers were sonicated in 20 ml sterile PBS-Tween solution (1x, 0.15%) for 10 minutes to remove epiphytic microbes. Flower parts were removed with sterile tweezers and the solution was centrifuged at 3000 rpm for 10 minutes to pellet the microbial cells. Both root and flower samples were stored at -80°C until DNA extraction. Aboveground wet biomass was assessed before plant material was dried at 50°C for at least 7 days to determine aboveground dry weight.

Bumblebees were euthanized by freezing at -20°C and frozen bees were dissected under sterile conditions to extract the gut. Gut samples were frozen at -80°C until extraction. Vole and earthworm feces were stored at -20°C.

**DNA extraction, library preparation and 16S rRNA gene amplicon sequencing**

To analyze the microbial composition, DNA of root, earthworm feces, vole feces and soil samples were extracted using the *ZymoBIOMICS DNA Miniprep* kit (Zymo Research Europe, Germany), whereas the *ZymoBIOMICS DNA Microprep* kit was used to extract DNA of flower and bumblebee gut samples. All samples were extracted following the manufacturer's instructions and subjected to an initial bead beating step using ceramic beads provided by the kit to mechanically lyse bacterial cells. To break open the roots for the endosphere samples five steel beads (2.4 mm, Bio-Budget Technologies, Germany) were additionally added to each sample. Bead beating was carried out using a SpeedMill PLUS (Analytik Jena, Germany) running two 3-minute cycles of bead-beating interrupted by a 3 - minute break.

To assess the bacterial composition, the hypervariable V4 region of the 16S rRNA gene was amplified in a two-step polymerase-chain reaction (PCR) using the primers 515 F (5'-GTGCCAGCMGCCGCGGTAA-3') and 806 R (5'-GGACTACHVGGGTWTCTAAT-3') [4]. To avoid amplifying high amounts of chloroplast and mitochondrial DNA in plant (endosphere and flower) and bumblebee gut samples, 10µM peptide nucleic acid (PNA)-DNA clamps (mPNA and pPNA in plant samples and mPNAs in bumblebee samples) were added to the first PCR reaction [5]. The PCR regime for these samples included one more annealing step for 30 seconds at 78°C after denaturation within the 30 cycles. PCR success for all samples was confirmed by agarose gel electrophoresis. Appended adapters for forward (CS1-515F) and reverse (CS2-806R) primers were used in order to construct the library using Standard Bio Tools chemistry (Access Array System for Illumina Sequencing Systems, Standard BioTools, USA) [6-8]. The second PCR reaction was performed to attach individual barcodes and adapters used for illumina sequencing. To account for potential sequencing artifices arising from the DNA extraction, a positive control mock community (Zymo Research Europe, Germany), as well as extraction blanks and PCR-negative controls, were also included as a part of the in-house Illumina Miseq sequencing run at the Institute of Evolutionary Ecology and Conservation Genomics at Ulm University in Germany.

**Bioinformatic sequence processing**

Sequence processing was conducted on Qiime2 (version 2022.8) using the DADA2 pipeline which incorporates primer removal, denoising, chimera removal and merging of paired-end reads [9,10]. Given the high number of samples and the samples being sequenced on two different runs, the denoising steps were conducted for each run separately. Files of the representative sequences, metadata files and the dada2 output table were merged before taxonomical assignment. For the first run forward reads were trimmed at 30 and 240 bp and reverse reads were trimmed at 30 and 215bp. For the second run, forward reads were trimmed at 30 and 240bp, reverse reads were trimmed at 30 and 220bp. For taxonomical assignment the SILVA 138-99 database [11] using the q2-feature classifier classify-sklearn (version 0.24.1) [12] in QIIME2 was used and run on the combined dataset of both sequencing runs. A phylogenetic tree was constructed after all ASVs were aligned with mafft [13] (via q2-alignment) using fasttree [14] (via q2-phylogeny). Using the software Dendroscope (version 3.8.4)[15], the unrooted phylogenetic tree was re-rooted using an uncultured archaeal sequence (accession number KT433146.1). The rooted phylogenetic tree and the biom table, both generated in Qiime2, together with the metadata file were imported into R using the phyloseq package [16] and a phyloseq object was created to conduct all statistical analysis using the R language [17] within the R Studio interface (version 4.3.2, Posit team (2023). The data processing of the phyloseq object in R precluded filtering of unassigned taxa as well as taxa assigned to chloroplast and mitochondria on family, class or order level and non-bacterial sequences. It was followed by filtering of samples by retaining the samples with non-zero reads.

**Table S1: Sample size distribution of the study.** Number of samples included in the microbiome analyses in relation to the seven different trophic compartments and fertilization regimes. Samples marked with an asterisk (*) represent pools of fecal samples of one species due to low volume of material collected at one sampling site. All other samples represent individual non-pooled samples.

| | Control | Biogas digestate | Cow/horse manure | Pig slurry | total |
|---|---|---|---|---|---|
| **Soil** | 24 | 24 | 20 | 24 | 92 |
| **Rhizosphere** (*Trifolium pratense*) | 24 | 24 | 20 | 24 | 92 |
| **Endosphere** (*Trifolium pratense*) | 24 | 24 | 20 | 24 | 92 |
| **Earthworm** | 6* | 6* | 4* | 6* | 22* |
| **Vole** (*Microtus Arvalis, Arvicola amphibious)* | 5 | 16 | 25 | 3 | 49 |
| **Flower** (*Trifolium pratense*) | 24 | 24 | 20 | 24 | 92 |
| **Bumblebee** *(Bombus lapidarius)* | 18 | 13 | 16 | 4 | 51 |
| **Pig Slurry** | 0 | 0 | 0 | 4 | 4 |
| **Total** | 125 | 131 | 125 | 109 | 494 |

**Table S2: Statistical results assessing the differences in alpha diversity (Shannon index) between the four different fertilization regimes within the seven different trophic compartments.** Indicated are p-values of ANOVAs and post-hoc Tukey's HSD pairwise comparisons. Significant values are marked in bold.

| Comparison | Soil | Rhizosphere | Endosphere | Vole | Earthworm | Flower | Bee |
|---|---|---|---|---|---|---|---|
| Overall | **0.029** | **0.001** | 0.145 | **0.001** | 0.107 | 0.124 | 0.473 |
| Control-pig | 0.075 | **0.001** | - | 0.091 | - | - | - |
| Control-biogas | 0.981 | 0.987 | - | **0.001** | - | - | - |
| Control-cow | 0.959 | 0.060 | - | **0.003** | - | - | - |
| Pig-biogas | **0.029** | **0.001** | - | 0.881 | - | - | - |
| Pig-cow | 0.262 | **0.049** | - | 0.999 | - | - | - |
| Biogas-cow | 0.826 | **0.026** | - | 0.549 | - | - | - |

**Table S3: Statistical results assessing differences in beta diversity (Bray-Curtis distances) between the four different fertilization regimes within the seven different trophic compartments.** P-values are given for PERMANOVA and pairwise PERMANOVA comparisons. Significant values are marked in bold.

| Comparison | All | Soil | Rhizo-sphere | Endo-sphere | Vole | Earth-worm | Flower | Bee |
|---|---|---|---|---|---|---|---|---|
| Overall | **0.001** | **0.001** | **0.001** | **0.001** | **0.001** | **0.005** | **0.001** | **0.001** |
| Control-pig | **0.001** | **0.001** | **0.001** | **0.001** | 0.656 | **0.021** | **0.001** | 0.229 |
| Control-biogas | **0.001** | **0.001** | **0.001** | **0.001** | **0.001** | 0.299 | **0.001** | **0.023** |
| Control-cow | **0.001** | **0.001** | **0.001** | **0.002** | **0.001** | **0.005** | **0.001** | 0.144 |
| Pig-biogas | **0.001** | **0.005** | **0.001** | **0.007** | **0.003** | 0.134 | **0.002** | **0.005** |
| Pig-cow | **0.001** | **0.005** | **0.001** | **0.001** | **0.005** | 0.121 | **0.002** | 0.37 |
| Biogas-cow | **0.003** | 0.153 | **0.009** | **0.01** | **0.002** | 0.052 | **0.015** | **0.001** |

**Table S4: Set of interactions to decipher exchange of microbes across the trophic chain.**

Compartments soil, rhizosphere or endosphere were considered as potential source while key target compartments were endosphere, flowers, earthworms and voles.

| | **Source** | | | |
|---|---|---|---|---|
| | Soil | Rhizosphere | Endosphere | Flower |
| **Sink** | Rhizosphere | Endosphere | Earthworm | Bumblebee |
| | Earthworm | Earthworm | Vole | |
| | Vole | Vole | | |
| | Flower | | | |

**Table S5: Statistical results assessing differences in beta diversity (Bray-Curtis distances) between the seven different trophic compartments across all fertilization regimes using the refined dataset containing only resilient and responding genera**. Indicated are p-values of Pairwise PERMANOVA comparisons. Significant values are marked in bold.

| | Soil | Rhizosphere | Endosphere | Earthworms | Voles | Flowers |
|---|---|---|---|---|---|---|
| Rhizosphere | **0.001** | | | | | |
| Endosphere | **0.001** | **0.001** | | | | |
| Earthworm | **0.001** | **0.001** | **0.001** | | | |
| Vole | **0.001** | **0.001** | **0.001** | **0.001** | | |
| Flower | **0.001** | **0.001** | **0.001** | **0.001** | **0.001** | |
| Bumblebee | **0.001** | **0.001** | **0.001** | **0.001** | **0.001** | **0.001** |

**Table S6: Statistical results assessing differences in beta diversity (Bray-Curtis) between the four different fertilization regimes across all trophic compartments using the refined dataset containing only resilient and responding genera.** Indicated are p-values of Pairwise PERMANOVA comparisons. Significant values are marked in bold.

| | Control | Biogas digestate | Cow/horse manure |
|---|---|---|---|
| Biogas digestate | **0.001** | | |
| Cow/horse manure | **0.001** | **0.007** | |
| Pig slurry | **0.001** | **0.001** | **0.001** |

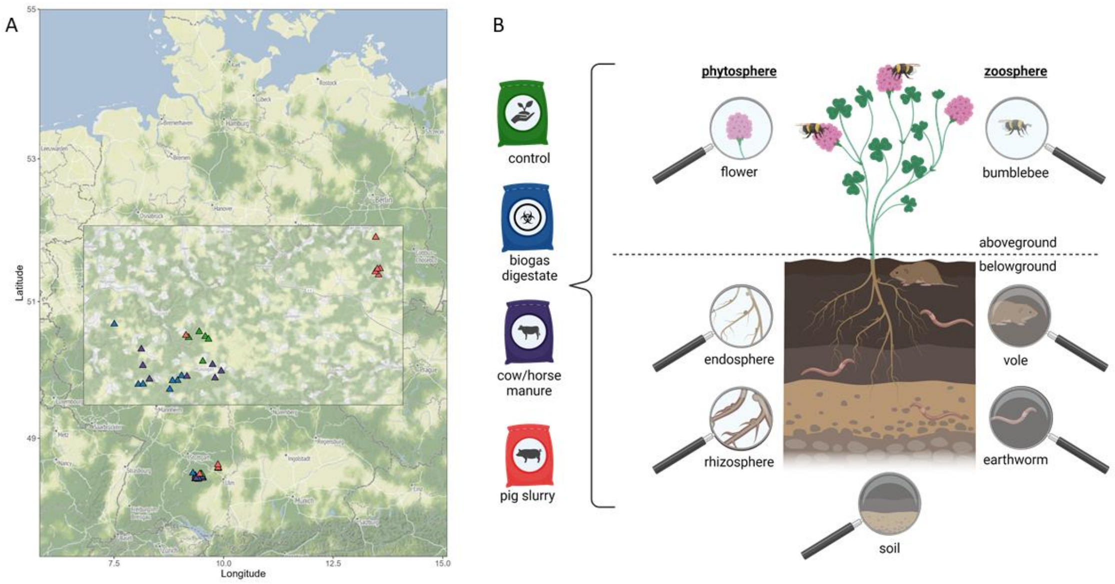


**Figure S1**: **Study design (A)** Map of sampling sites located on the Swabian Alb in Baden-Württemberg, Germany, subjected to four different fertilization regimes (green: minimally fertilized control sites, blue: sites fertilized with biogas digestate, purple: sites fertilized with cow/horse manure, pink: sites fertilized with pig slurry). Map was created using R. (B) Study set up with seven different compartments of the trophic chain: soil, rhizosphere, root endosphere and flower (*Trifolium pratense)*, earthworm, vole (*Microtus arvalis, Arvicola amphibious*) and bumblebee (B*ombus lapidarius*) as well as the four different fertilization regimes: minimally fertilized control, biogas digestate fertilization, cow/horse manure fertilization and pig slurry fertilization. Compartments of the trophic chain can be grouped either in aboveground (flower and bumblebee) and belowground (soil, rhizosphere, endosphere, vole and earthworm) or into phytosphere compartments (rhizosphere, endosphere and flower) and zoosphere compartments (earthworm, vole, bumblebee).

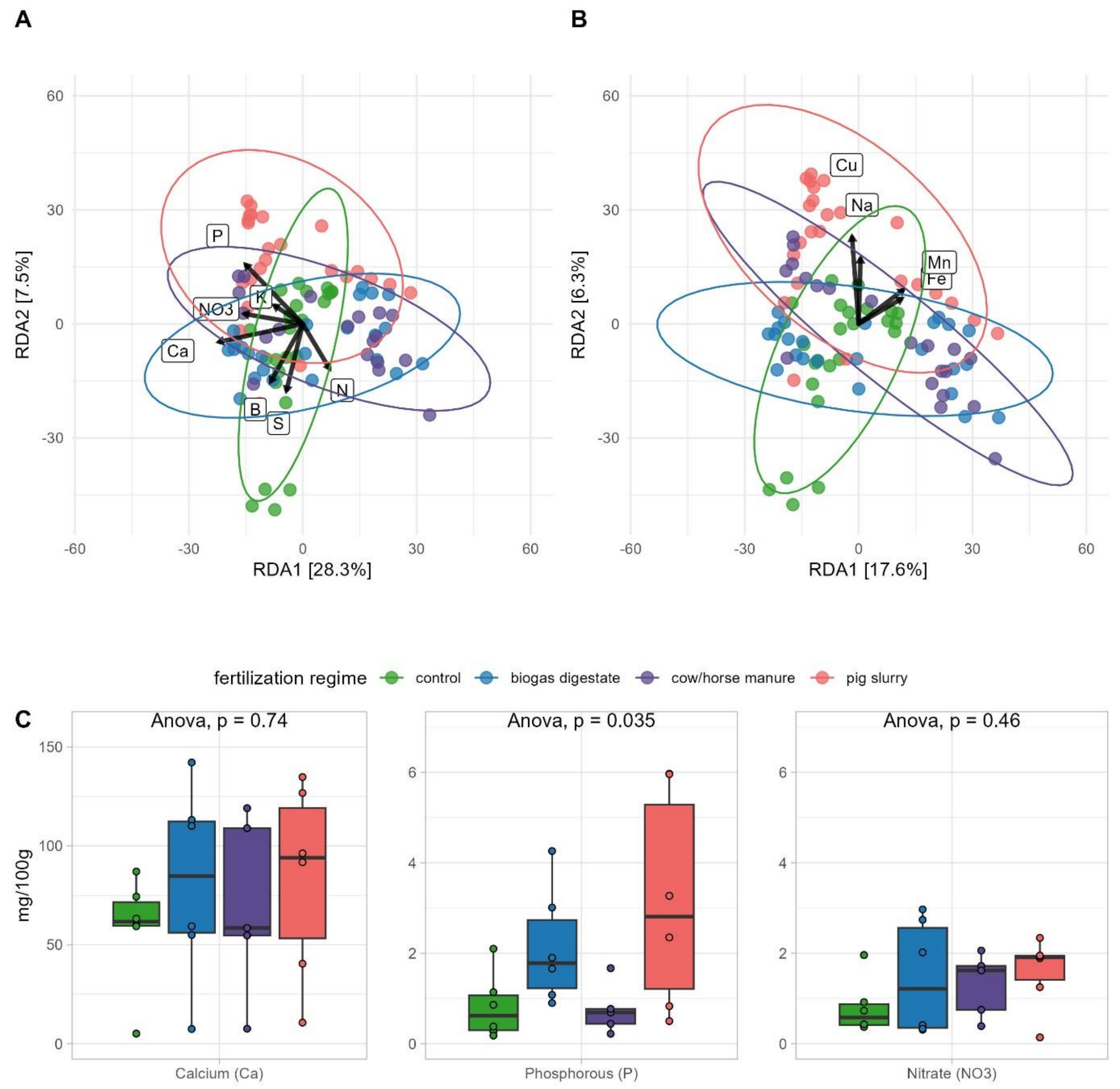


**Figure S2: Soil nutrient content analysis.** Redundancy analysis (RDA) modelling the effect of different soil (A) macronutrients and (B) micronutrients on Bray-Curtis distances of soil microbial communities and nutrient contents (C) of the 3 most important nutrients driving the soil microbial communities. Arrows in A and B indicate the direction of the different soil nutrients associated with the respective soil microbial communities, labels on the arrows indicate the respective soil nutrient. The colors represent the fertilization regime in which the soil samples were collected (green: minimally fertilized control sites, blue: biogas digestate fertilized sites, purple: cow/horse manure fertilized sites, pink: pig slurry fertilized sites). (C) ANOVA was used to assess differences between fertilization regimes in the soil content of the three nutrients explaining the most variation in microbial communities (Ca, P and $NO_3^-$).

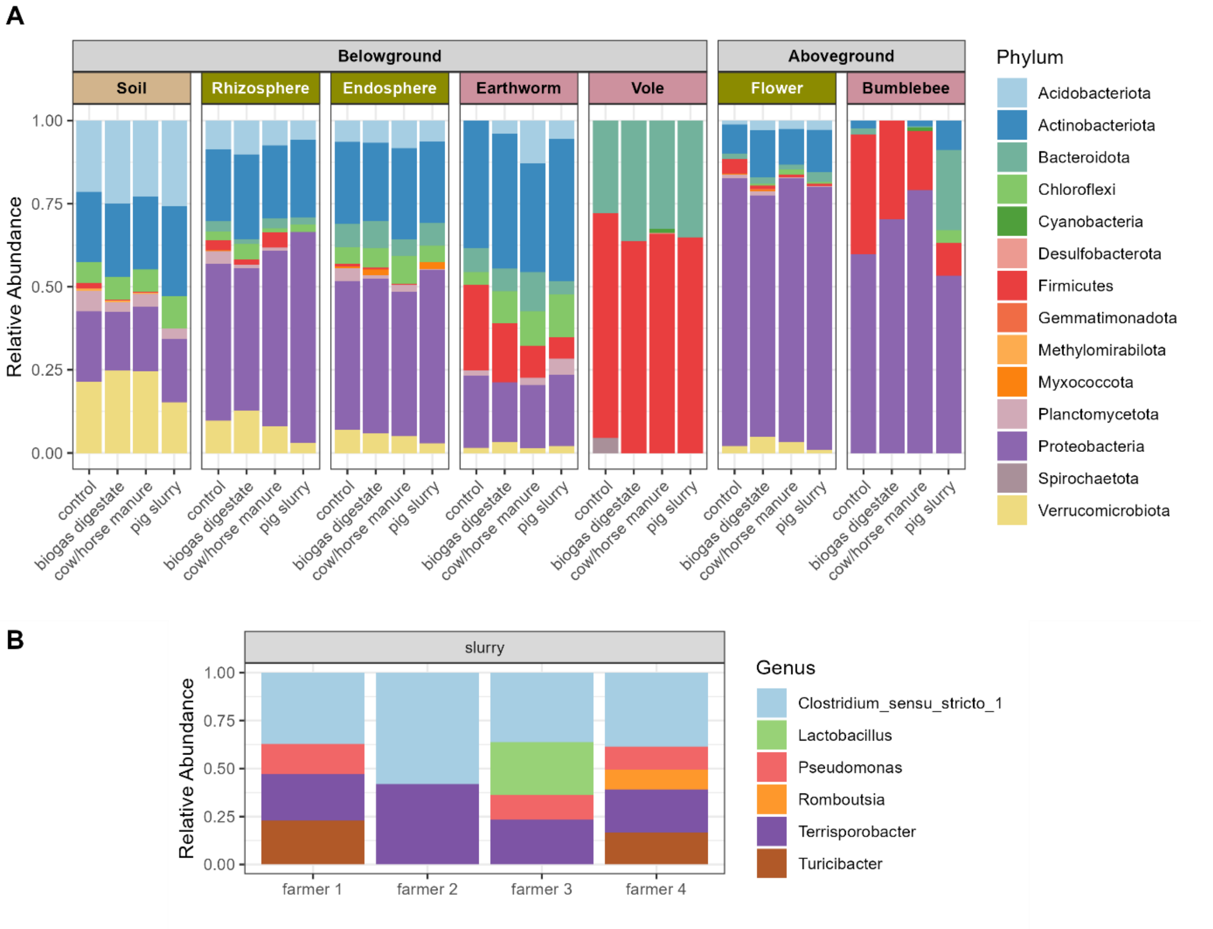


**Figure S3**: **Bacterial composition of the different trophic compartments and pig slurry** (A) Bacterial composition of all trophic compartments indicating phyla with a minimum abundance of 5%. Corresponding to Fig. 2, data was pooled for plotting based on the fertilization regime within each compartment. Every fertilization regime contains up to 6 plots with 4 samples each, i.e. each bar represents a pool of up to 24 samples (see Table S1 for sample sizes). (B) Bacterial composition of pig slurry samples obtained from farm sites before application to the field indicating genera with a minimum abundance of 5%. Data represents individual samples.

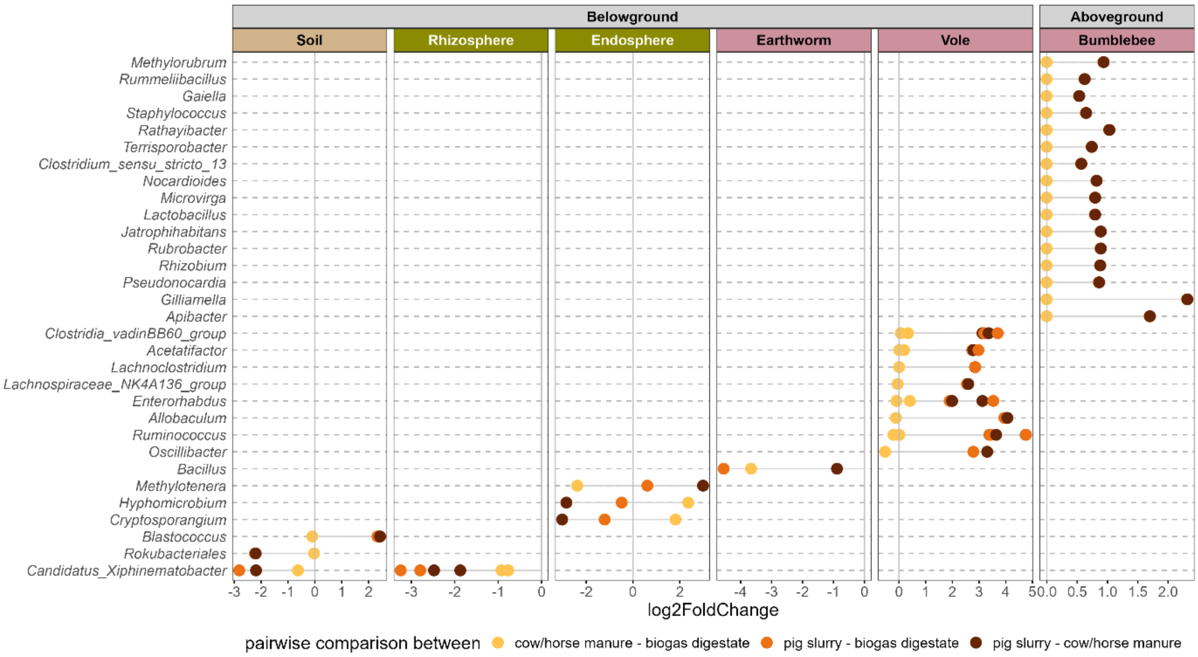


**Figure S4: Bacterial genera whose abundance changes in response to fertilization in different trophic compartments.** Pairwise comparisons were made between cow and horse manure/biogas digestate (yellow), between sites fertilized with pig slurry/biogas digestate (orange) and between sites fertilized with pig slurry/cow and horse manure (brown) using ANCOMBC2. Flowers did not have any identifiable genera being significantly different for the tested comparisons and are hence not shown. Positive values of log2fold change represent genera that are more abundant in the second partner in the comparison; negative values represent genera that are more abundant in the first partner in the comparison.

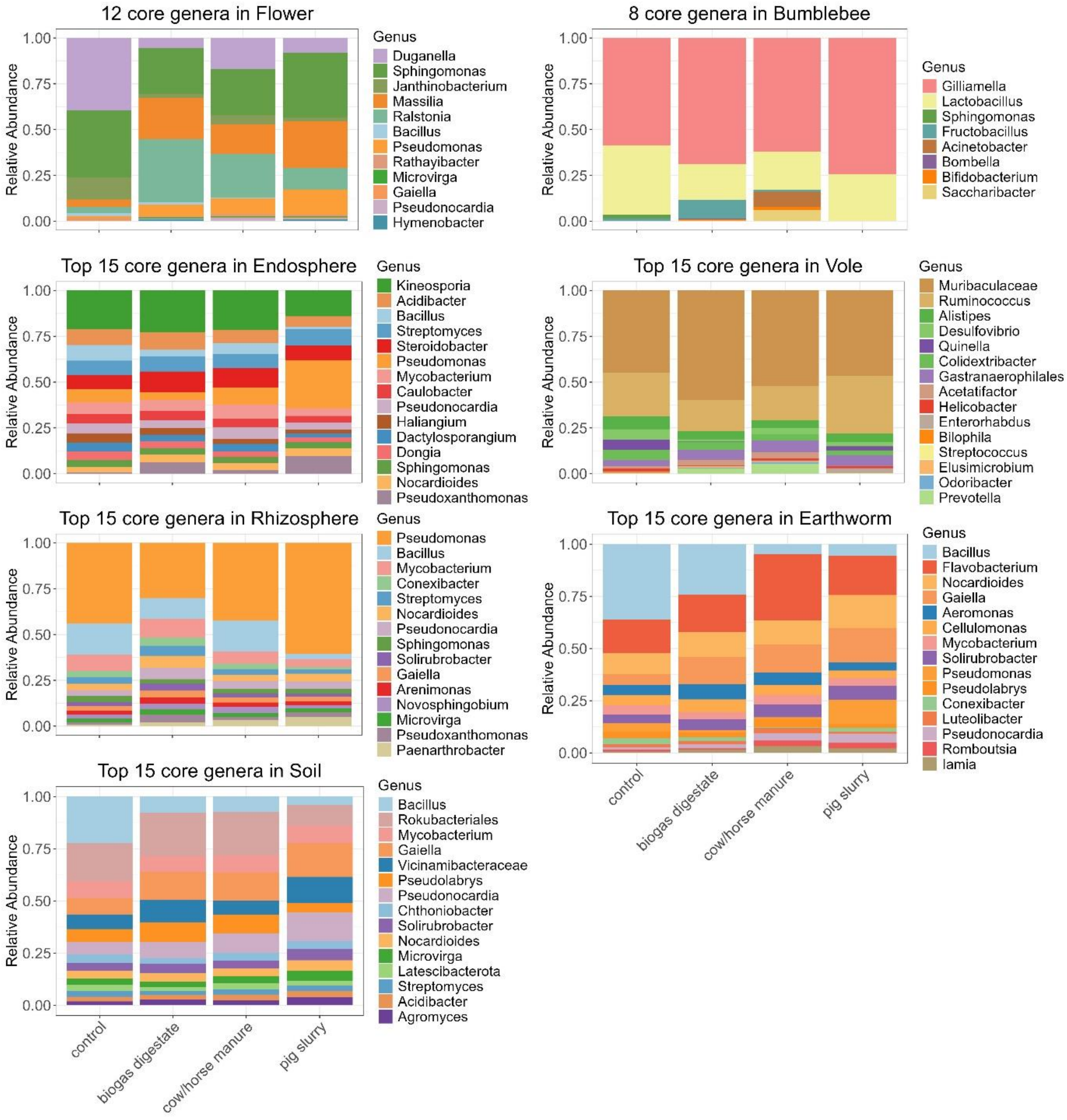


**Figure S5: Composition bar charts of the core genera in the different compartments along the trophic chain.** Indicated are the 15 most abundant core genera in soil, rhizosphere, endosphere, earthworms and voles, as well as the 12 core genera in flowers and 8 core genera in bumblebees in samples collected from sites subjected to the four different fertilization regimes (minimally fertilized control sites, biogas digestate fertilized sites, cow/horse manure fertilized sites and pig slurry fertilized sites).

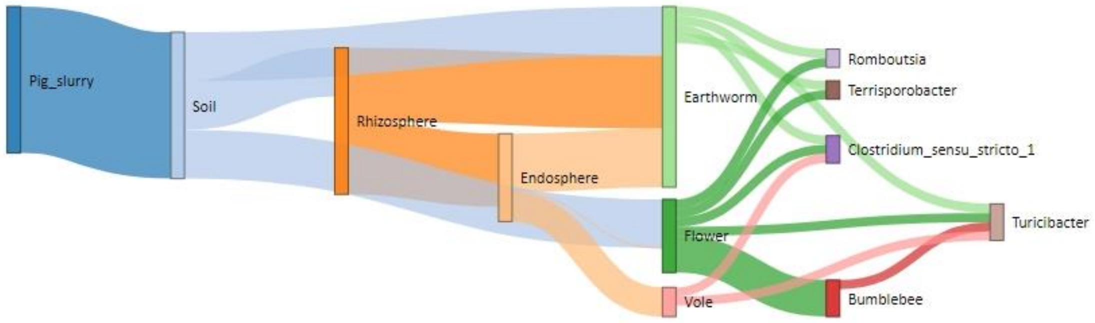


**Figure S6**: **Sourcetracker analysis showing the proportions of bacterial genera from pig slurry shared between compartments along the trophic chain.** Pig slurry was the primary source to all subsequent compartments, i.e. soil, rhizosphere, endosphere, while the main target compartments were earthworms, voles and flowers.

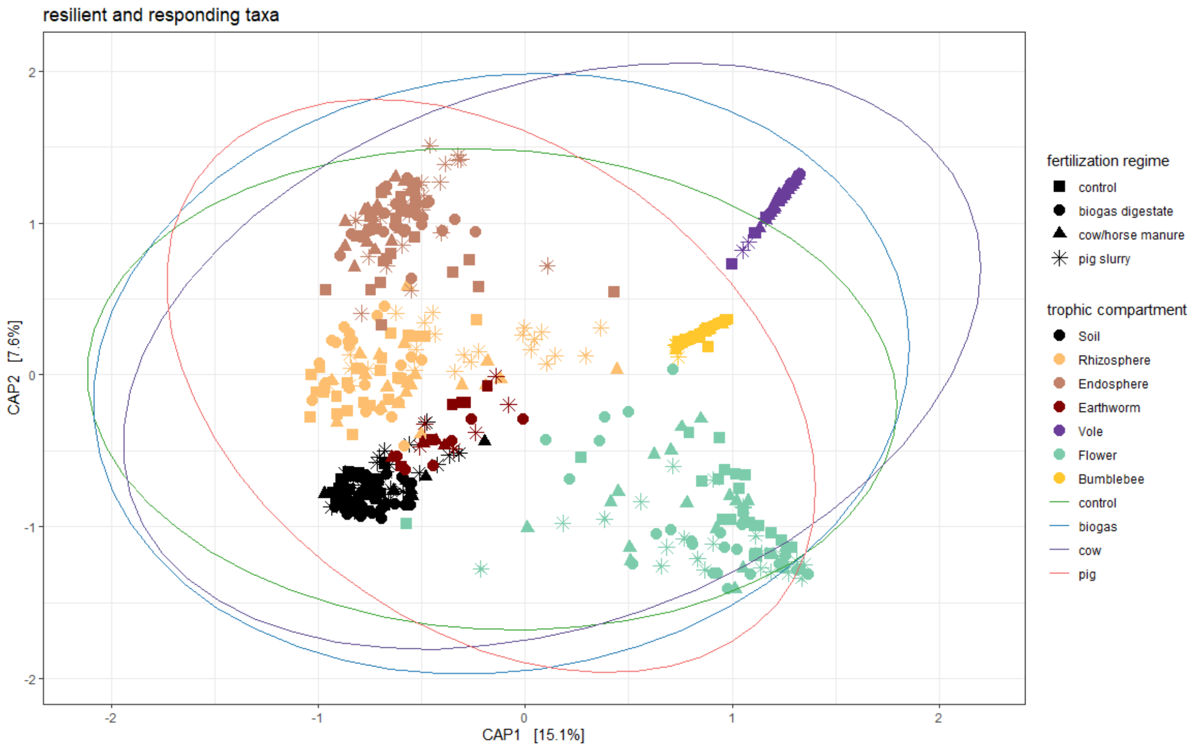


**Figure S7: Two-dimensional illustration of the refined dataset containing resilient and responder taxa identified by CAP analysis**. The different trophic compartments are indicated by colors and the fertilization regimes by different shapes.
